## Supplementary Material for "Gain-of-function mutations amplify cytotoxic FAM111 protease activity in human genetic disorders"

**Supplementary Figure 1.**

**Sequence conservation and structural modeling of FAM111 protease domains**

**a**. Sequence alignment of the serine protease domain-containing portions of FAM111A proteins from different mammals. Patient-associated mutations in human FAM111A (circles) and catalytic triad residues in the protease domain (stars) are indicated. **b**. Alignment of the serine protease domains in human FAM111A, human FAM111B and *E. coli* DegS. **c**. Crystal structure of monomeric *E. coli* DegS protease (PDB ID: 2R3U). Zoomed-in view shows position of catalytic triad residues (yellow). **d**. Homology-based protease domain model of human FAM111A (residues 371-555; teal). Zoomed-in view shows catalytic triad residues (yellow). Residues mutated in human disease (red) are indicated. **e**. Homology-based protease domain model of human FAM111B (residues 471-664; blue). Zoomed-in view shows catalytic triad residues (yellow). Residues mutated in human disease (red) are indicated. **f**. Overlay of the DegS, FAM111A and FAM111B protease domains (c-e).

**Supplementary Figure 2.**

**Elevated FAM111A proteolytic activity impairs DNA replication and triggers replication stress**

**a.** Representative images of U2OS/GFP-FAM111A cell lines that were treated or not with DOX, fixed and stained with PCNA antibody. Mutation of the PIP box abrogates FAM111A localization to PCNA-positive DNA replication foci. **b**. DNA replication rates in HCT116/GFP-FAM111A WT cells treated with DOX, pulse-labeled with EdU and stained with DAPI were analyzed by quantifying EdU signal intensity in S phase cells using QIBC ($n$>500 cells per condition; red bars, mean (A.U., arbitrary units)). **c**. As in (a), except that cells were co-stained with PCNA and RPA2 antibodies. **d**. Quantification of data in (c) (grey bars, average; *n*>2000 cells per condition). Scale bars, 10 µm.

Data (**a-d**) are representative of three independent experiments with similar outcomes.

**Supplementary Figure 3.**

**FAM111A and FAM111B are dispensable for DNA replication and cell proliferation**

**a**. Immunoblot analysis of U2OS cells transfected with indicated siRNAs. **b**. Cells in (a) were pulse-labeled with EdU, fixed and stained with DAPI. Cells were then subjected to QIBC analysis for quantification of EdU and DAPI signal intensities (*n*>2000 cells per condition). Proportion of cells in G1 (blue), S (red) and G2/M (yellow) phases is indicated. **c**. Immunoblot analysis of U2OS cells and derivative lines with targeted knockout of *FAM111A* and/or *FAM111B*. **d**. Proliferation of cell lines in (c) was determined by quantification of Resazurin incorporation (mean±s.d.; *n*=3 independent experiments). **e**. As in (b), but using U2OS cell lines with targeted knockout of *FAM111A* and/or *FAM111B*. **f**. DNA fiber analysis of indicated U2OS cell lines labeled with CldU (25 min; red) followed by IdU (25 min; green). Fork speeds were calculated as length of labeled track divided by pulse time (black bars, median±s.d.; at least 200 fibers analysed per condition). A representative fiber image is shown.

Data (**a-f**) are representative of three independent experiments with similar outcomes.

**Supplementary Figure 4.**

**Characterization of the interaction between FAM111A and RFC1**

**a**. Validation of FAM111A interactors identified by mass spectrometry (Fig. 1h). U2OS/GFP-FAM111A WT cells treated or not with DOX were subjected to GFP immunoprecipitation (IP) followed by immunoblotting with indicated antibodies. **b**. U2OS cells transfected with constructs expressing indicated FLAG-tagged RFC1 fragments were subjected to FLAG IP followed by immunoblotting with indicated antibodies. **c**. U2OS cells transfected with non-targeting control (CTRL) or FAM111A siRNAs were pre-extracted, fixed and stained with FAM111A antibody. **d**. U2OS cells labeled with EdU were pre-extracted, fixed and stained with indicated antibodies. Endogenous FAM111A localizes to nucleoli in G1 and G2 phase (EdU-negative) cells and relocates to nuclear foci in S phase (EdU-positive) cells. **e**. U2OS cells transfected with indicated siRNAs were pre-extracted, fixed and stained with FAM111A and RFC1 antibodies. **f**. Immunoblot analysis of U2OS/GFP-FAM111A cell lines treated with DOX. Scale bars, 10 μm.

Data (**a-f**) are representative of three independent experiments with similar outcomes.

**Supplementary Figure 5.**

**FAM111A proteolytic activity triggers apoptosis in a Caspase-dependent but p53-independent manner**

**a**. U2OS/GFP-FAM111A cell lines treated or not with DOX were fixed, stained with propidium iodide (PI) and analyzed by flow cytometry. Proportion of cells with sub-G1 DNA content (red gate) is indicated. Approx. 30,000 cells were analyzed per condition. **b**. U2OS/GFP-FAM111A WT cells treated with DOX and/or pan-Caspase inhibitor Z-VAD-FMK for 24 h as indicated were fixed and stained with γ-H2AX antibody. Cells were then subjected to QIBC analysis of γ-H2AX signal intensity ($n$>2000 cells per condition; red bars, mean (A.U., arbitrary units)). **c**. U2OS/GFP-FAM111A WT cells transfected with indicated siRNAs for 48 h were treated or not with DOX for an additional 24 h were immunoblotted with indicated antibodies. **d**. Immunoblot analysis of HCT116/GFP-FAM111A WT cells treated with DOX for the indicated times. **e**. Cells treated as in (b) were labeled with EdU, fixed and stained with DAPI. Cells were then subjected to QIBC analysis of EdU signal intensity (*n*>1000 cells per condition). **f**. U2OS/GFP-FAM111A WT cells treated with DOX for the indicated times were fixed and stained with γ-H2AX antibody. Cells were then subjected to QIBC analysis of γ-H2AX signal intensity (red bars, mean; *n*>2000 cells per condition). **g**. U2OS cell lines treated with DOX and labeled with EdU were fixed and stained with γ-H2AX antibody. Cells were then subjected to QIBC analysis of γ-H2AX signal intensity (red bars, mean; *n*>2000 cells per condition). Cells were classified as being in G1, S and G2/M phases based on DNA content and EdU signal intensity. **h**. U2OS/GFP-FAM111A cell lines treated with DOX were pulse-labeled with EU, stained with DAPI and analyzed by QIBC (*n*>2000 cells per condition).

Data (**a**-**h**) are representative of three independent experiments with similar outcomes.

**Supplementary Figure 6.**

**Patient-associated mutations hyperactivate FAM111A protease activity to exacerbate its adverse impact on cellular fitness**

**a**. Immunoblot analysis of parental U2OS cells (-) or derivative stable cell lines conditionally expressing GFP-FAM111 WT at different levels. **b**. Immunoblot analysis of parental U2OS cells (-) or derivative stable cell lines expressing WT or patient-associated GFP-FAM111A alleles. **c**. Cells in (b) were pulse-labeled with EdU, fixed and stained with DAPI. Cells were then subjected to QIBC analysis for quantification of EdU and DAPI signal intensities (*n*>2000 cells per condition; A.U., arbitrary units). **d**. Quantification of data in (c) (red bars, mean). **e**. Representative images of U2OS/GFP-FAM111A cell lines that were treated or not with DOX for the indicated times, fixed and co-stained with PCNA and RPA2 antibodies. Scale bar, 10 μm. **f**. Quantification of data in (e) (grey bars, average; *n*>2000 cells per condition). **g**. DOX-treated U2OS/GFP-FAM111A cell lines were stained with γ-H2AX antibody and analyzed for γ-H2AX signal intensity by QIBC (red bars, mean; *n*>2000 cells per condition). **h**. As in (g), except that cells were stained with RFC1 antibody, pre-extracted and fixed, and stained with DAPI. RFC1 and DAPI signal intensities were analyzed by QIBC (red bars, mean; *n*>2000 cells per condition). **i**. As in (g), except that cells were stained with RPB1 antibody and analyzed for RPB1 signal intensity by QIBC (red bars, mean; *n*>2000 cells per condition). **j**. U2OS cell lines conditionally expressing untagged ectopic FAM111A alleles were treated or not with DOX for 24 h, labeled with EdU, fixed and stained with DAPI. Cells were then subjected to QIBC analysis for quantification of EdU and DAPI signal intensities (*n*>2000 cells per condition). **k**. Immunoblot analysis of stable U2OS/FAM111A cell lines transfected or not with FAM111A siRNA targeting the 3'UTR, and subsequently treated or not with DOX to express ectopic untagged FAM111A alleles.

Data (**a-k**) are representative of three independent experiments with similar outcomes.

**Supplementary Figure 7.**

**Characterization of purified recombinant FAM111 proteins**

**a**. Recombinant human FLAG-tagged FAM111A proteins purified from yeast were analyzed by Coomassie staining. **b**. Immunoblot analysis of recombinant FLAG-FAM111A proteins in (a). **c**. Recombinant human FLAG-tagged FAM111B proteins purified from yeast were analyzed by Coomassie staining.

Data (**a-c**) are representative of two independent experiments with similar outcomes.

**Supplementary Figure 8.**

**Elevated FAM111A protease activity disrupts microtubule organization**

**a-c**. Representative images of U2OS/GFP-FAM111A cell lines that were treated or not with DOX for the indicated times, then left untreated or exposed to nocodazole for 1 h, and fixed and stained

with α-tubulin antibody. Note that cells with elevated FAM111A protease activity display accumulation of nuclear α-tubulin. **d**. Immunoblot analysis of U2OS/GFP-FAM111A D528G cells treated with DOX and Z-VAD-FMK for 8 h, as indicated. **e**. U2OS cells were transfected with the indicated siRNAs, fixed 4 days later and stained with crystal violet. **f**. Immunoblot analysis of U2OS/GFP-FAM111A WT and mutants induced with DOX for 16 h. All scale bars, 10 μm.
Data are representative of three (**a**-**c**) and two (**d**-**f**) independent experiments with similar outcomes.

**Supplementary Figure 9.**

**Patient-associated FAM111B mutants trigger DNA replication suppression, apoptosis onset and microtubule network disruption in a protease-dependent manner**

**a**. Sequence alignment of the serine protease domain-containing portions of FAM111B proteins from different mammals. Patient-associated mutations in human FAM111B (circles) and catalytic triad residues in the protease domain (stars) are indicated. **b**. Immunoblot analysis of U2OS/GFP-FAM111B cell lines treated with DOX for 24 h to induce expression of WT or disease-associated GFP-FAM111B alleles. **c**. Cells treated as in (b) were fixed, stained with γ-H2AX antibody and DAPI, and γ-H2AX signal intensity was analyzed by QIBC (red bars, mean; *n*>2000 cells per condition; A.U., arbitrary units). **d**. Cells treated as in (b) were labeled with EdU, fixed and stained with DAPI. EdU signal intensity was analyzed by QIBC (red bars, mean; *n*>2000 cells per condition). **e**. Representative images of U2OS/GFP-FAM111B cell lines that were treated or not with DOX, then left untreated or exposed to nocodazole for 1 h, and fixed and stained with α-tubulin antibody.
Data (**b**-**e**) are representative of three independent experiments with similar outcomes.

**Supplementary Figure 10.**

**Patient-associated FAM111B mutants interact with and displace RFC1 and RPB1 from chromatin**

**a**. U2OS/GFP-FAM111B S628N cells left untreated or exposed to DOX for 16 h were pulse-labeled with EdU, pre-extracted and fixed. Cells were then stained with indicated antibodies and DAPI, and analyzed by QIBC (*n*>2000 cells per condition; A.U., arbitrary units). **b**. U2OS cells transfected with indicated RFC subunit expression plasmids were subjected to FLAG IP and immunoblotted with indicated antibodies. **c**. U2OS/GFP-FAM111A or FAM111B WT cells treated with DOX were subjected to GFP IP followed by immunoblotting with indicated antibodies. **d**. Representative images of U2OS/GFP-FAM111B cell lines that were treated or not with DOX, fixed and stained with PCNA and RPA2 antibodies. Scale bar, 10 µm. **e**. Quantification of data in (d) (grey bars, average; *n*>2000 cells per condition). **f**. U2OS/GFP-FAM111B cell lines treated with DOX were pre-extracted, fixed and stained with RPB1 antibody. RPB1 signal intensity was analyzed by QIBC (*n*>2000 cells per condition). **g**. Quantification of data in (f) (red bars, mean; *n*>2000 cells per condition).

Data (**a-g**) are representative of three independent experiments with similar outcomes.

**Supplementary Figure 11.**

**DNA replication suppression and apoptosis induction by disease-associated FAM111A does not require FAM111B and *vice versa***

**a**. Immunoblot analysis of U2OS/GFP-FAM111A D528G cells transfected with control (CTRL) or FAM111B siRNAs, and subsequently left untreated or exposed to DOX. **b**. Cells treated as in (a) were pulse-labeled with EdU, stained with DAPI and analyzed for DAPI and EdU signal intensity using QIBC (*n*>2000 cells per condition; A.U., arbitrary units). **c**. Cells treated as in (a) were fixed, stained with γ-H2AX antibody and DAPI, and γ-H2AX signal intensity was analyzed by QIBC (red bars, mean; *n*>2000 cells per condition). **d**. Immunoblot analysis of U2OS/GFP-FAM111B S628N cells transfected with control (CTRL) or FAM111A siRNAs, and subsequently left untreated or exposed to DOX. **e**. Cells treated as in (d) were pulse-labeled with EdU, stained with DAPI and analyzed for DAPI and EdU signal intensity using QIBC (*n*>2000 cells per condition). **f**. Cells treated as in (d) were fixed, stained with γ-H2AX antibody and DAPI, and γ-H2AX signal intensity was analyzed by QIBC (red bars, mean; *n*>2000 cells per condition).

Data (**a**-**f**) are representative of three independent experiments with similar outcomes.

**Supplementary Figure 12.**

***FAM111A* and *FAM111B* mRNA expression levels in different human tissues**

Human Protein Atlas RNA-seq data on *FAM111A* and *FAM111B* transcript expression levels in 37 human tissues (pTPM, protein transcripts per million).

a

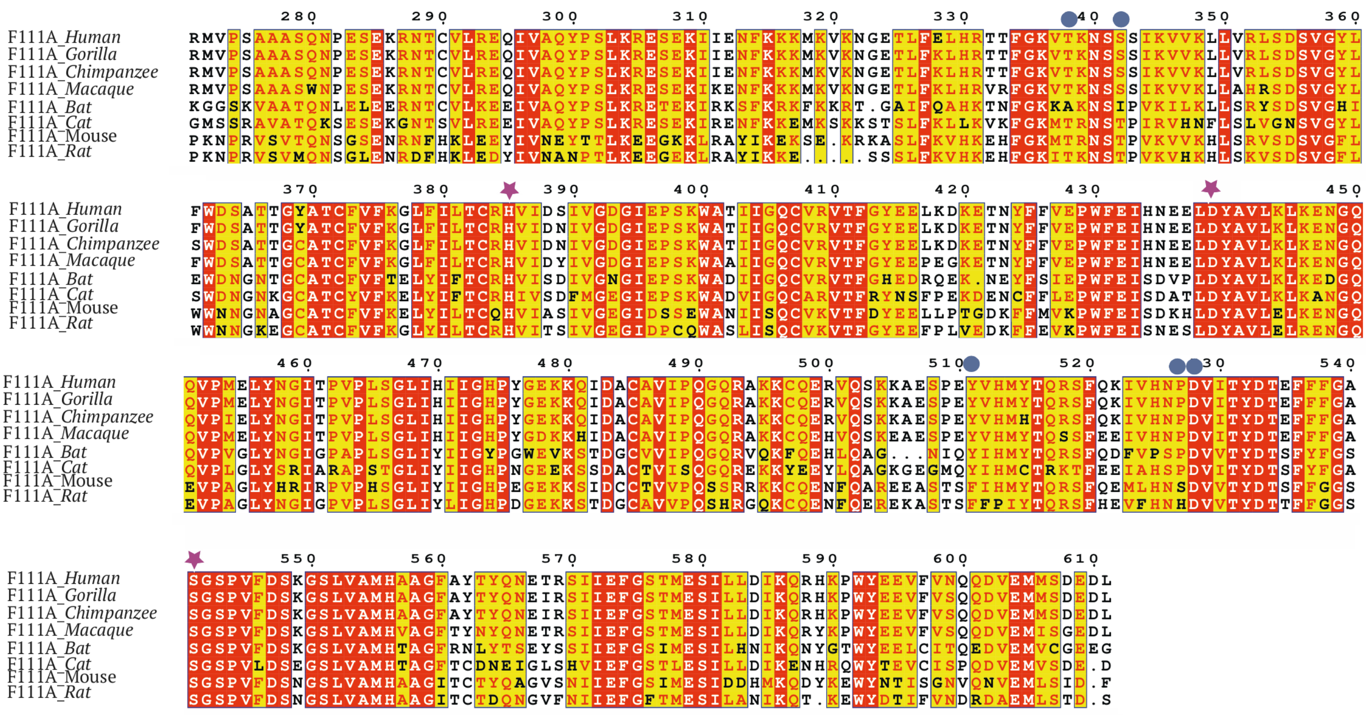
F111A_Human
F111A_Gorilla
F111A_Chimpanzee
F111A_Macaque
F111A_Bat
F111A_Cat
F111A_Mouse
F111A_Rat

b

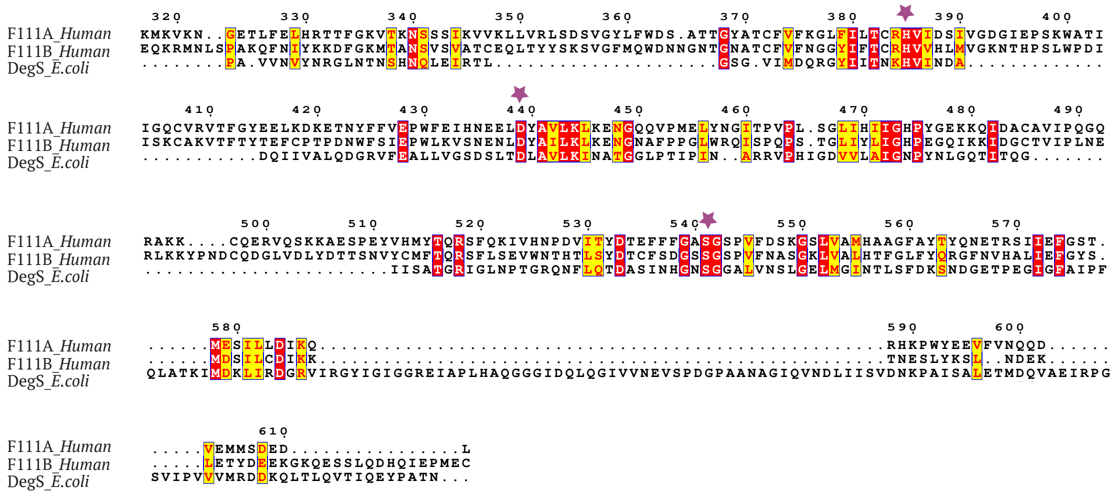
F111A_Human
F111B_Human
DegS_E.coli

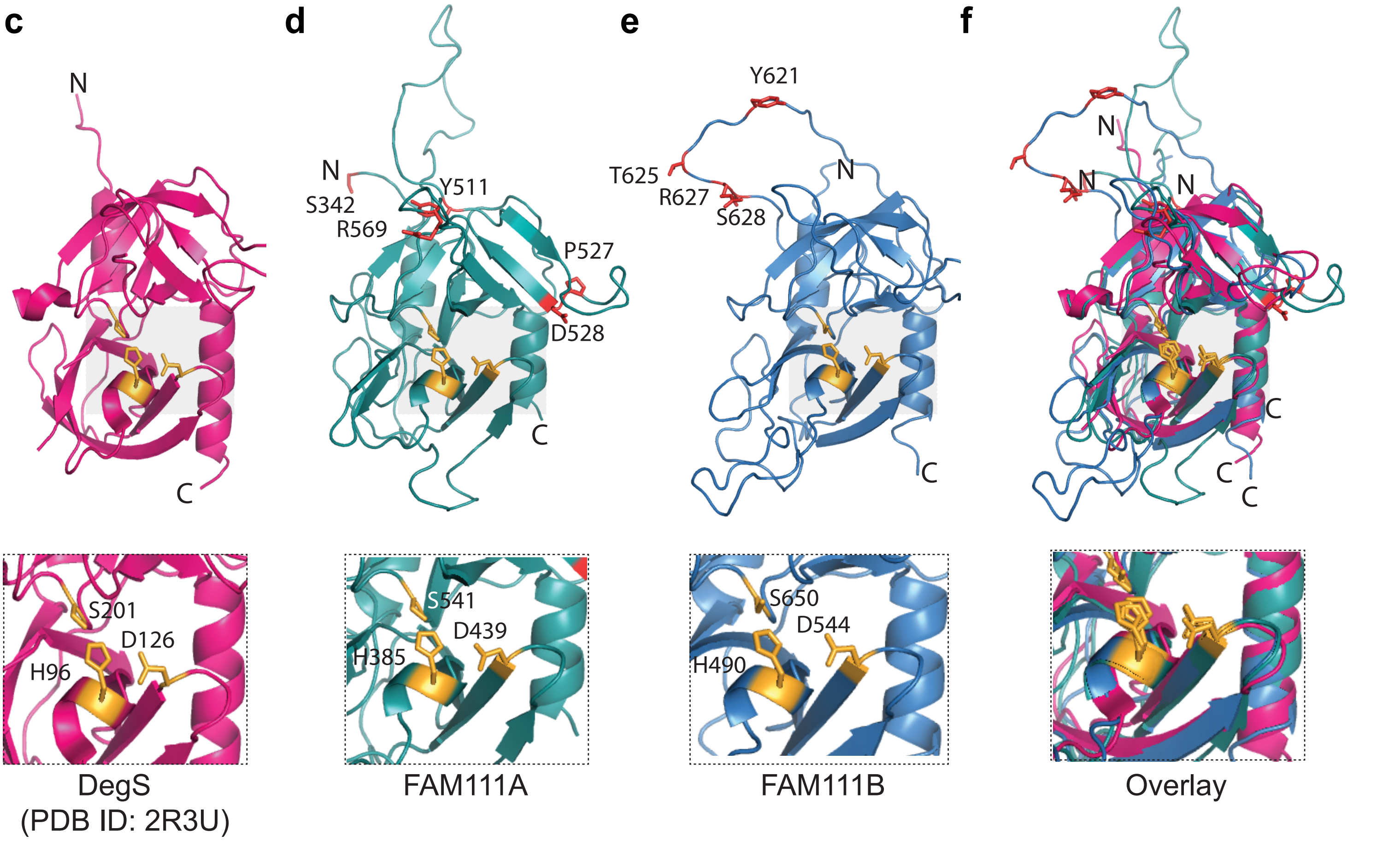
c
d
e
f
N
C
S342
Y511
R569
P527
D528
Y621
T625
R627
S628
S201
D126
H96
S541
D439
H385
S650
D544
H490
DegS
(PDB ID: 2R3U)
FAM111A
FAM111B
Overlay

Hoffmann et al., Supplementary Figure 2

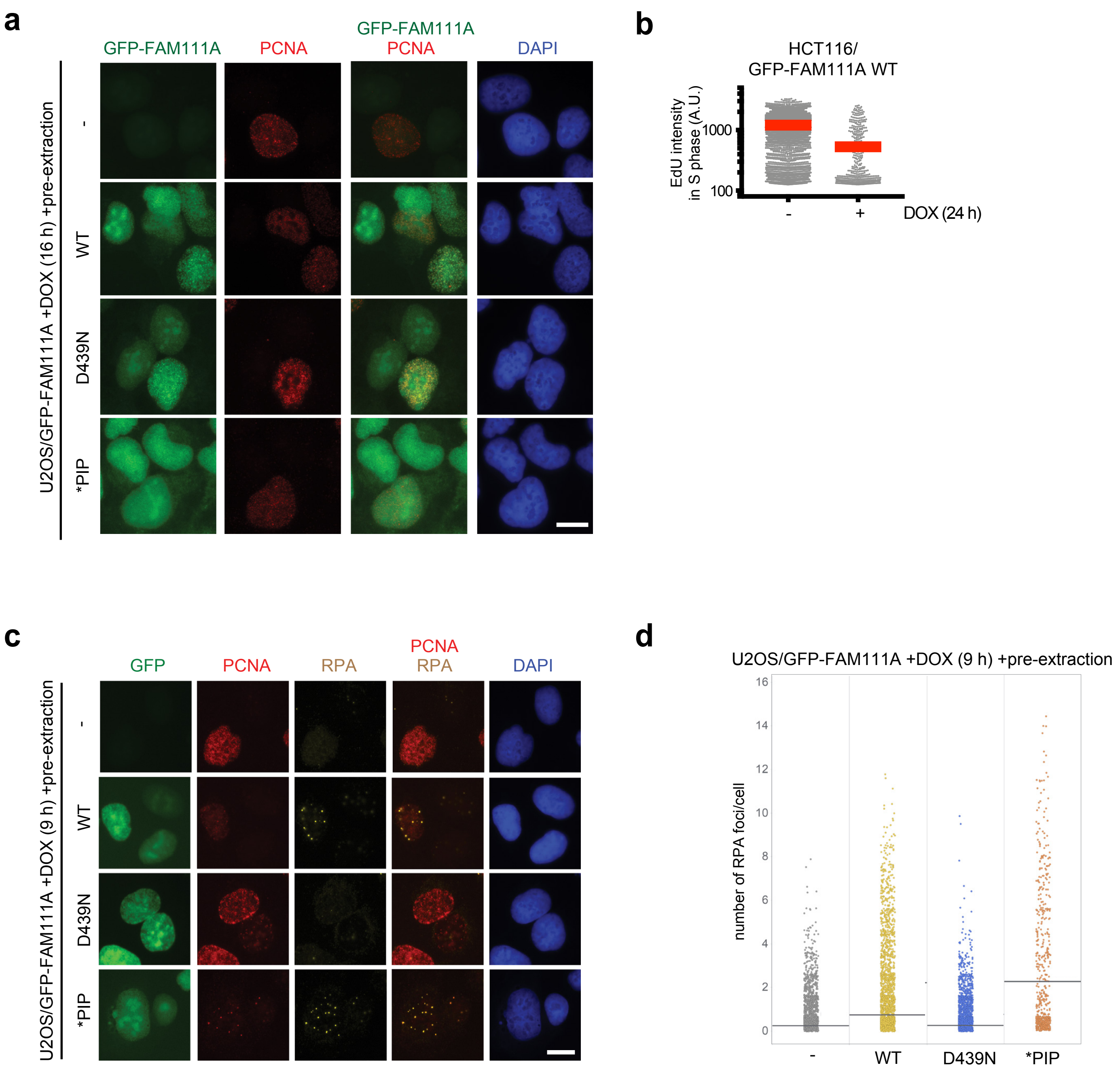

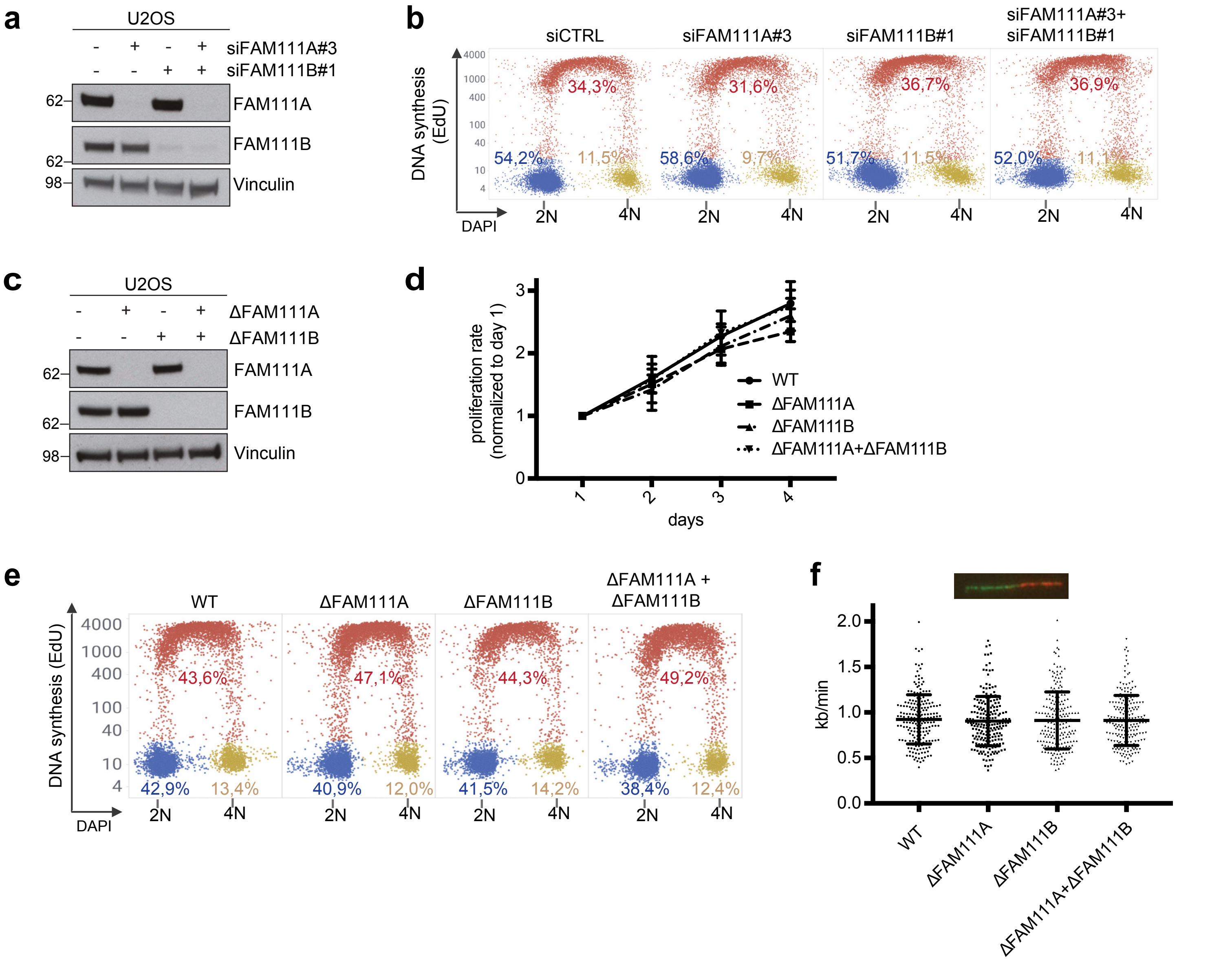
a
U2OS
siFAM111A#3
siFAM111B#1
FAM111A
FAM111B
Vinculin
b
siCTRL
siFAM111A#3
siFAM111B#1
siFAM111A#3+ siFAM111B#1
DNA synthesis (EdU)
DAPI
34,3%
31,6%
36,7%
36,9%
54,2%
11,5%
58,6%
9,7%
51,7%
11,5%
52,0%
11,1%
c
ΔFAM111A
ΔFAM111B
d
proliferation rate (normalized to day 1)
days
WT
ΔFAM111A
ΔFAM111B
ΔFAM111A+ΔFAM111B
e
WT
ΔFAM111A
ΔFAM111B
ΔFAM111A + ΔFAM111B
43,6%
47,1%
44,3%
49,2%
42,9%
13,4%
40,9%
12,0%
41,5%
14,2%
38,4%
12,4%
f
kb/min

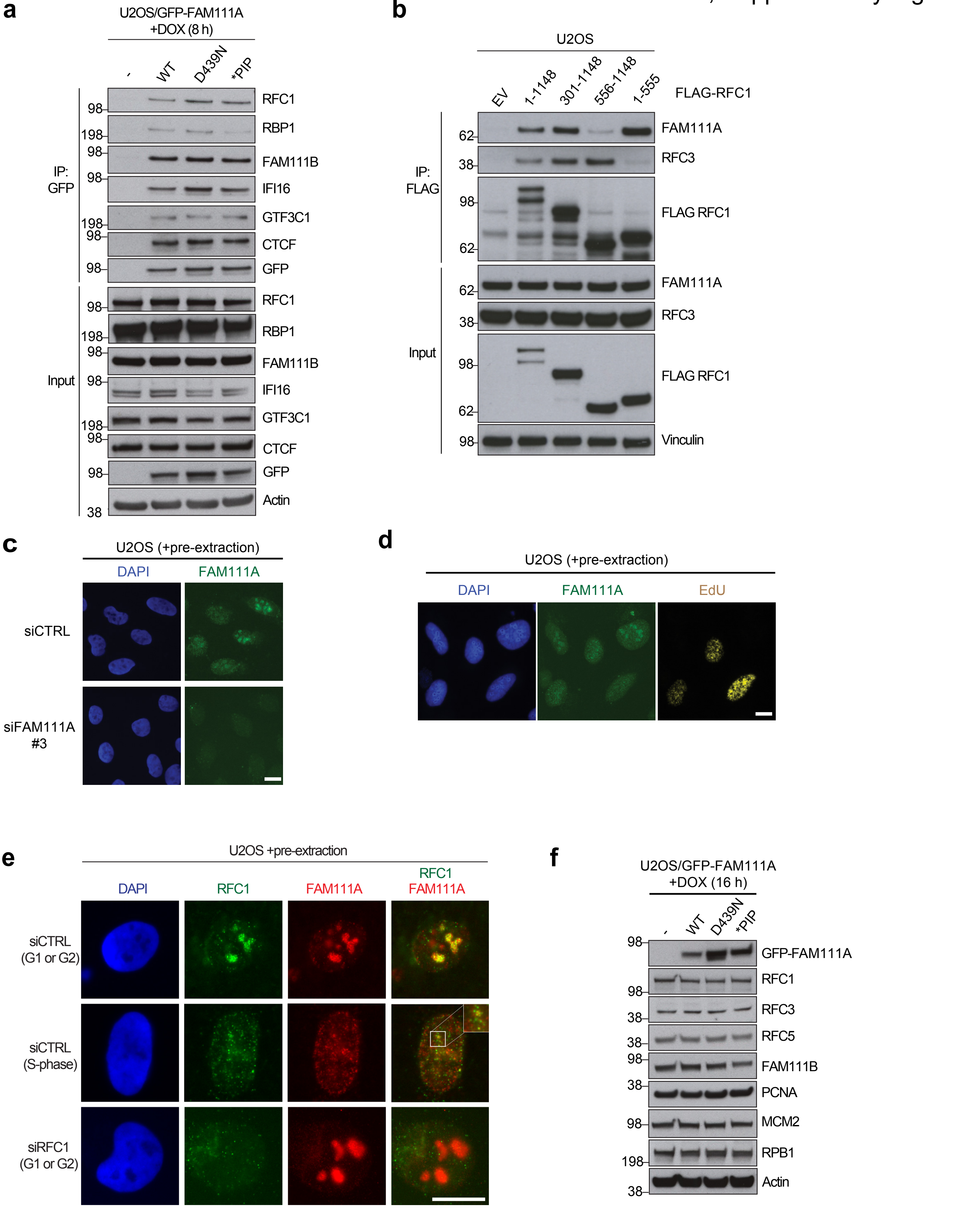
a
U2OS/GFP-FAM111A +DOX (8 h)
- WT D439N *PIP
IP: GFP
RFC1
RBP1
FAM111B
IFI16
GTF3C1
CTCF
GFP
Input
RFC1
RBP1
FAM111B
IFI16
GTF3C1
CTCF
GFP
Actin
98 198 98 98 198 98 98 98 198 98 98 198 98 98 38
b
U2OS
EV 1-1148 301-1148 556-1148 1-555
FLAG-RFC1
IP: FLAG
FAM111A
RFC3
FLAG RFC1
Input
FAM111A
RFC3
FLAG RFC1
Vinculin
62 38 98 62 62 38 98 62 98
c
U2OS (+pre-extraction)
DAPI
FAM111A
siCTRL
siFAM111A #3
d
U2OS (+pre-extraction)
DAPI
FAM111A
EdU
e
U2OS +pre-extraction
DAPI
RFC1
FAM111A
RFC1 FAM111A
siCTRL (G1 or G2)
siCTRL (S-phase)
siRFC1 (G1 or G2)
f
U2OS/GFP-FAM111A +DOX (16 h)
- WT D439N *PIP
GFP-FAM111A
RFC1
RFC3
RFC5
FAM111B
PCNA
MCM2
RPB1
Actin
98 98 38 38 98 38 98 198 38

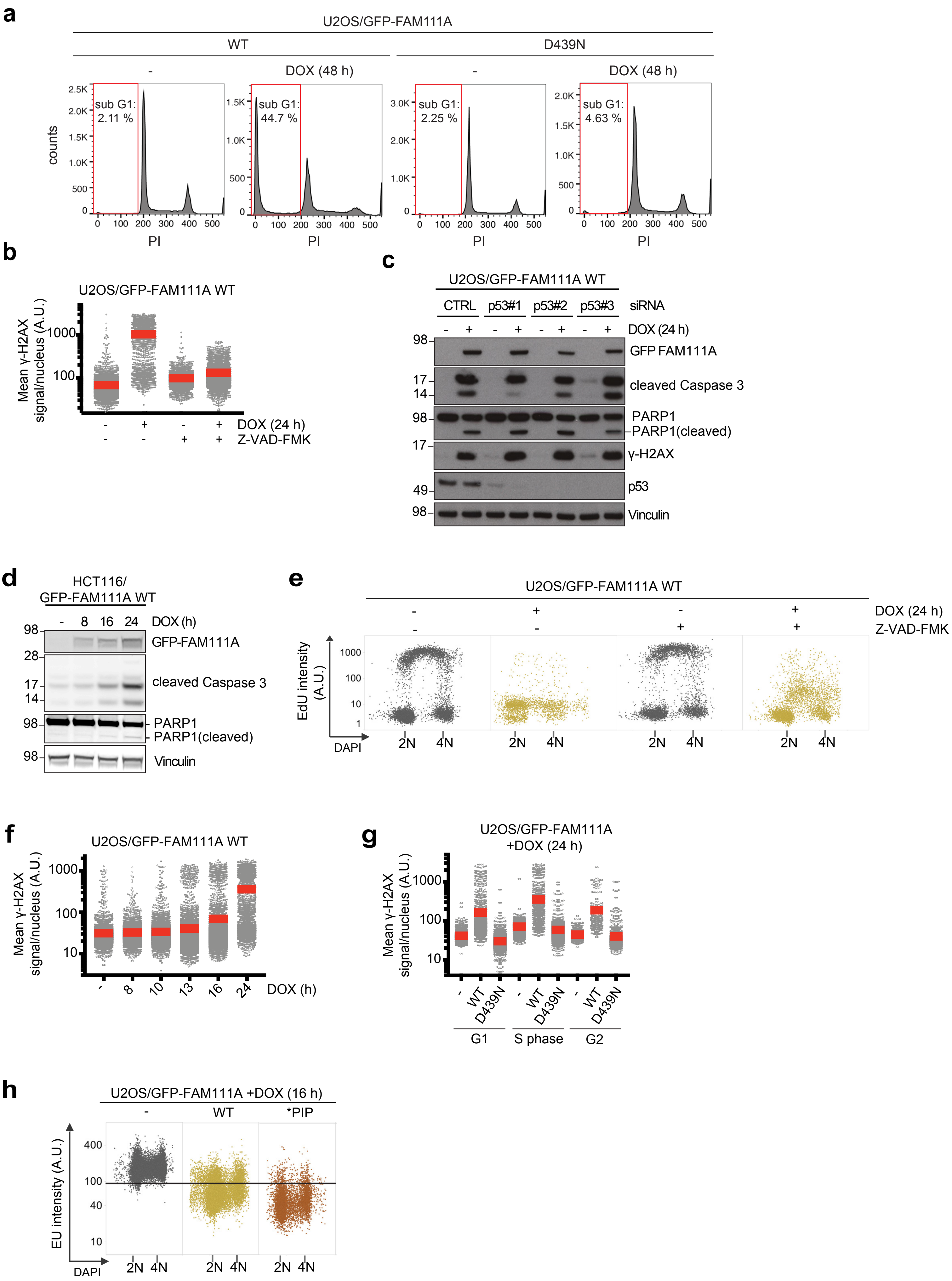
a
U2OS/GFP-FAM111A
WT
D439N
-
DOX (48 h)
sub G1: 2.11 %
sub G1: 44.7 %
sub G1: 2.25 %
sub G1: 4.63 %
counts
PI
b
U2OS/GFP-FAM111A WT
Mean γ-H2AX signal/nucleus (A.U.)
DOX (24 h)
Z-VAD-FMK
c
U2OS/GFP-FAM111A WT
CTRL
p53#1
p53#2
p53#3
siRNA
DOX (24 h)
GFP FAM111A
cleaved Caspase 3
PARP1
PARP1(cleaved)
γ-H2AX
p53
Vinculin
d
HCT116/ GFP-FAM111A WT
DOX (h)
GFP-FAM111A
cleaved Caspase 3
PARP1
PARP1(cleaved)
Vinculin
e
U2OS/GFP-FAM111A WT
DOX (24 h)
Z-VAD-FMK
EdU intensity (A.U.)
DAPI
2N
4N
f
U2OS/GFP-FAM111A WT
Mean γ-H2AX signal/nucleus (A.U.)
DOX (h)
g
U2OS/GFP-FAM111A +DOX (24 h)
Mean γ-H2AX signal/nucleus (A.U.)
WT
D439N
G1
S phase
G2
h
U2OS/GFP-FAM111A +DOX (16 h)
WT
*PIP
EU intensity (A.U.)
DAPI
2N
4N

Hoffmann et al., Supplementary Figure 6

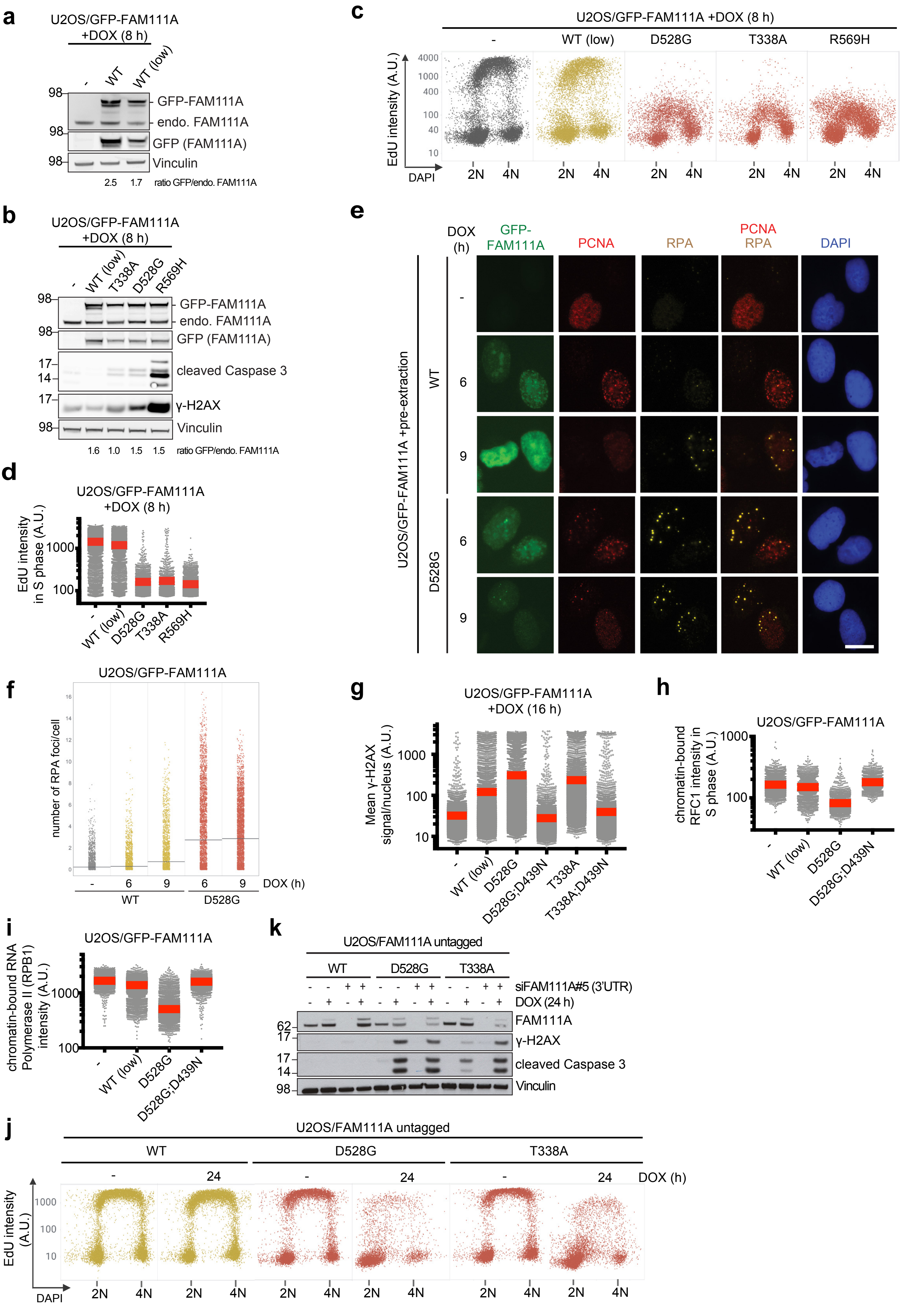

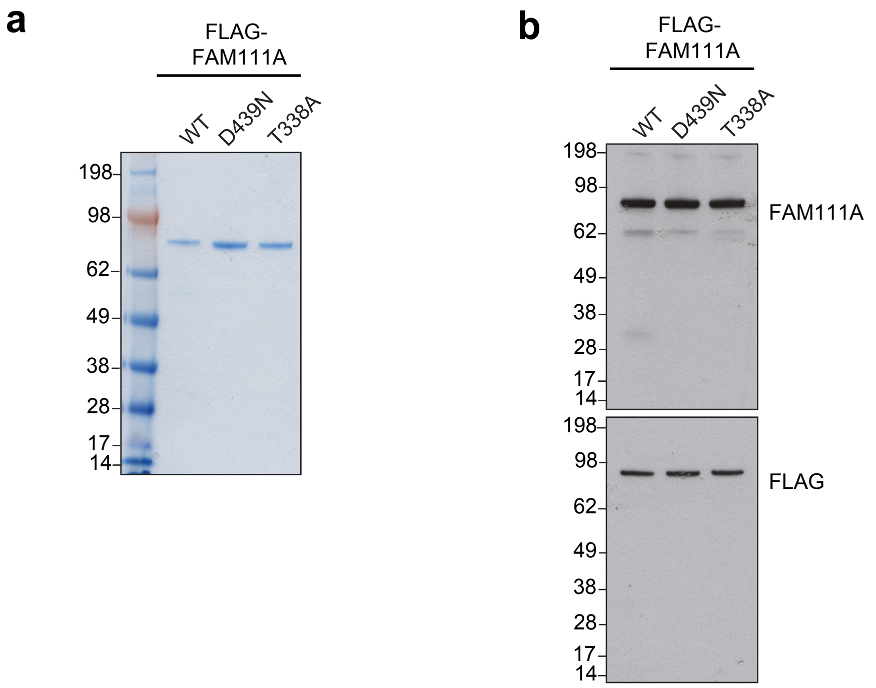
a
FLAG-
FAM111A
WT
D439N
T338A
198
98
62
49
38
28
17
14
b
FLAG-
FAM111A
WT
D439N
T338A
198
98
62
49
38
28
17
14
FAM111A
198
98
62
49
38
28
17
14
FLAG
c
FLAG-
FAM111B
WT
D544N
S628N
198
98
62
49
38
28
17
14

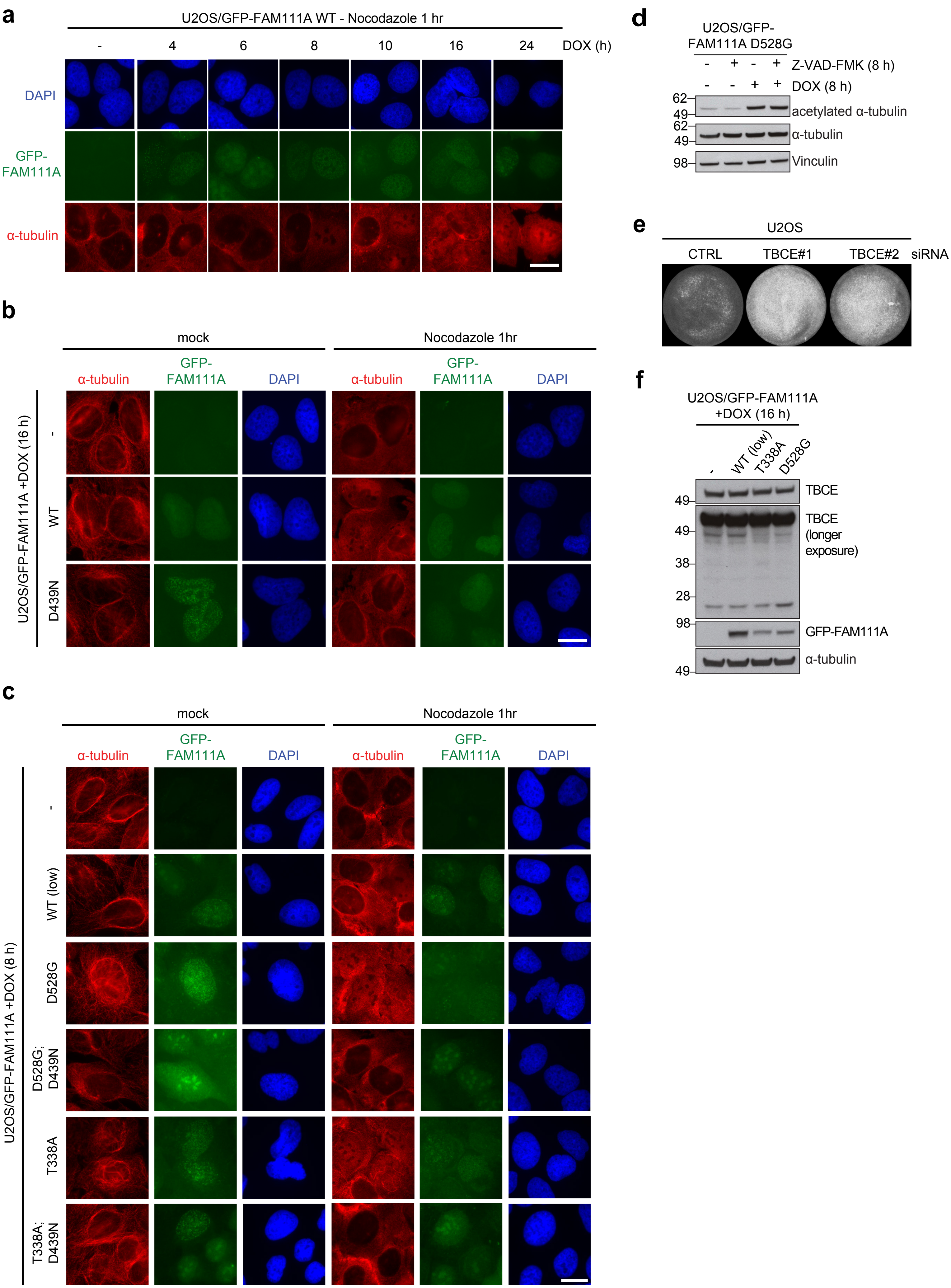
a
U2OS/GFP-FAM111A WT - Nocodazole 1 hr
- 4 6 8 10 16 24 DOX (h)
DAPI
GFP-FAM111A
α-tubulin
b
mock
Nocodazole 1hr
α-tubulin
GFP-FAM111A
DAPI
U2OS/GFP-FAM111A +DOX (16 h)
-
WT
D439N
c
mock
Nocodazole 1hr
α-tubulin
GFP-FAM111A
DAPI
U2OS/GFP-FAM111A +DOX (8 h)
-
WT (low)
D528G
D528G; D439N
T338A
T338A; D439N
d
U2OS/GFP-FAM111A D528G
- + - + Z-VAD-FMK (8 h)
- - + + DOX (8 h)
62
49
acetylated α-tubulin
62
49
α-tubulin
98
Vinculin
e
U2OS
CTRL
TBCE#1
TBCE#2
siRNA
f
U2OS/GFP-FAM111A +DOX (16 h)
-
WT (low)
T338A
D528G
49
TBCE
49
38
28
TBCE (longer exposure)
98
GFP-FAM111A
49
α-tubulin

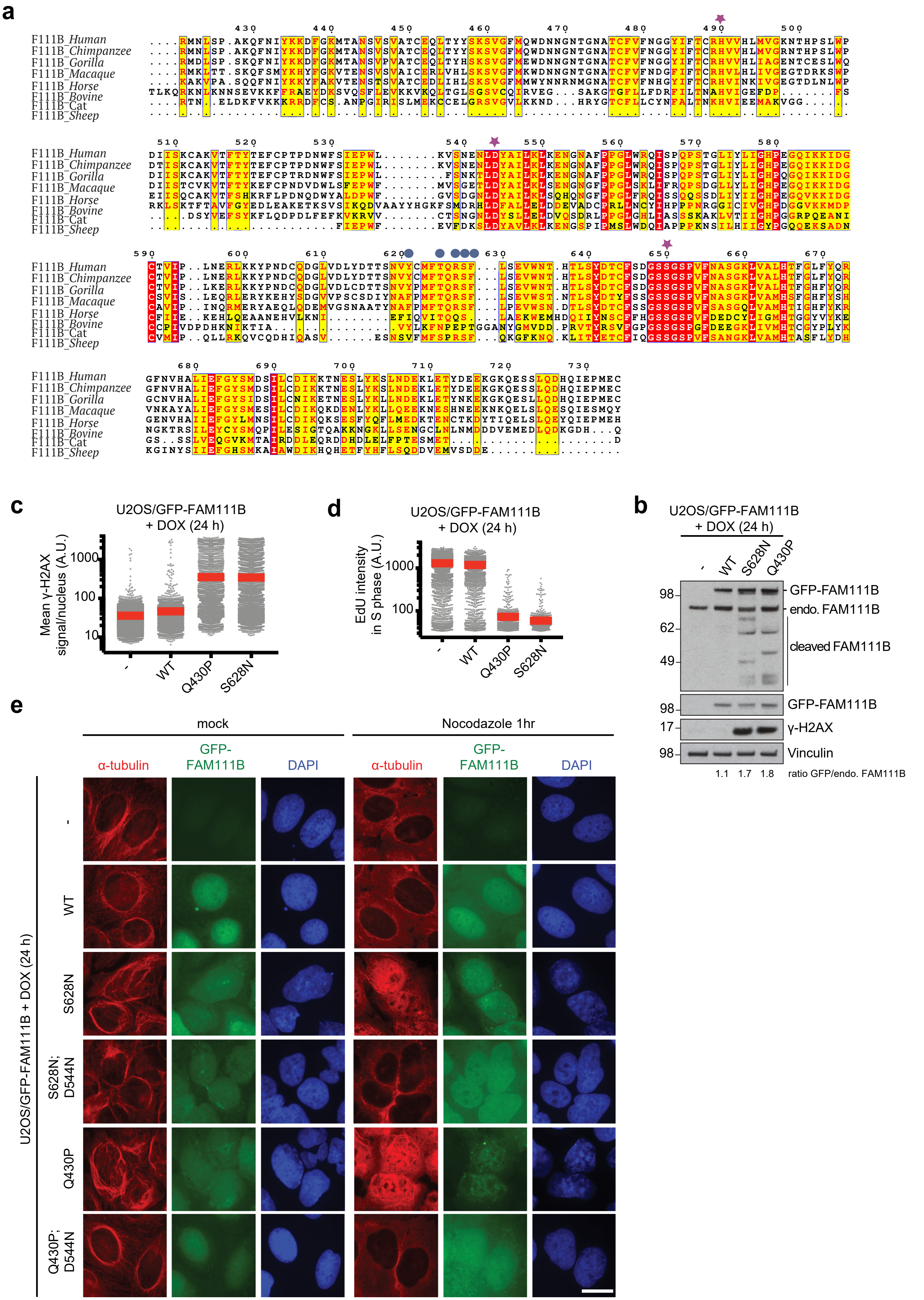
a
F111B_Human
F111B_Chimpanzee
F111B_Gorilla
F111B_Macaque
F111B_Horse
F111B_Bovine
F111B_Cat
F111B_Sheep
c
U2OS/GFP-FAM111B + DOX (24 h)
Mean γ-H2AX signal/nucleus (A.U.)
- WT Q430P S628N
d
U2OS/GFP-FAM111B + DOX (24 h)
EdU intensity in S phase (A.U.)
- WT Q430P S628N
b
U2OS/GFP-FAM111B + DOX (24 h)
- WT S628N Q430P
GFP-FAM111B
endo. FAM111B
cleaved FAM111B
γ-H2AX
Vinculin
1.1 1.7 1.8 ratio GFP/endo. FAM111B
e
mock
Nocodazole 1hr
α-tubulin GFP-FAM111B DAPI
U2OS/GFP-FAM111B + DOX (24 h)
- WT S628N S628N; D544N Q430P Q430P; D544N

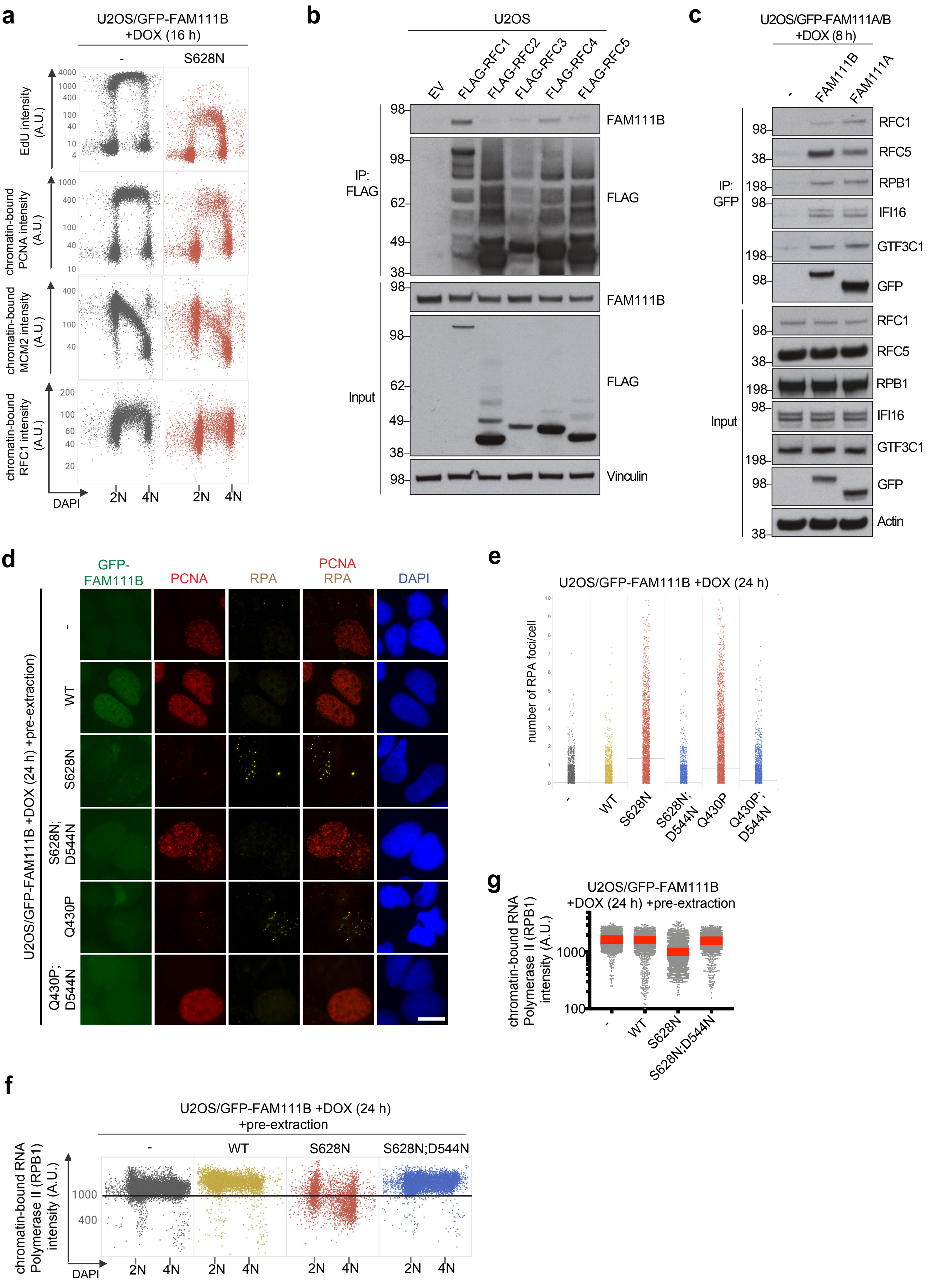

a
U2OS/GFP-FAM111B +DOX (16 h)
-
S628N
EdU intensity (A.U.)
chromatin-bound PCNA intensity (A.U.)
chromatin-bound MCM2 intensity (A.U.)
chromatin-bound RFC1 intensity (A.U.)
DAPI
2N 4N
b
U2OS
EV FLAG-RFC1 FLAG-RFC2 FLAG-RFC3 FLAG-RFC4 FLAG-RFC5
IP: FLAG
Input
FAM111B
FLAG
Vinculin
c
U2OS/GFP-FAM111A/B +DOX (8 h)
- FAM111B FAM111A
IP: GFP
Input
RFC1
RFC5
RPB1
IFI16
GTF3C1
GFP
Actin
d
GFP-FAM111B
PCNA
RPA
PCNA RPA
DAPI
U2OS/GFP-FAM111B +DOX (24 h) +pre-extraction
-
WT
S628N
S628N; D544N
Q430P
Q430P; D544N
e
U2OS/GFP-FAM111B +DOX (24 h)
number of RPA foci/cell
- WT S628N S628N; D544N Q430P Q430P; D544N
f
U2OS/GFP-FAM111B +DOX (24 h) +pre-extraction
- WT S628N S628N;D544N
chromatin-bound RNA Polymerase II (RPB1) intensity (A.U.)
DAPI
2N 4N
g
U2OS/GFP-FAM111B +DOX (24 h) +pre-extraction
chromatin-bound RNA Polymerase II (RPB1) intensity (A.U.)
- WT S628N S628N;D544N

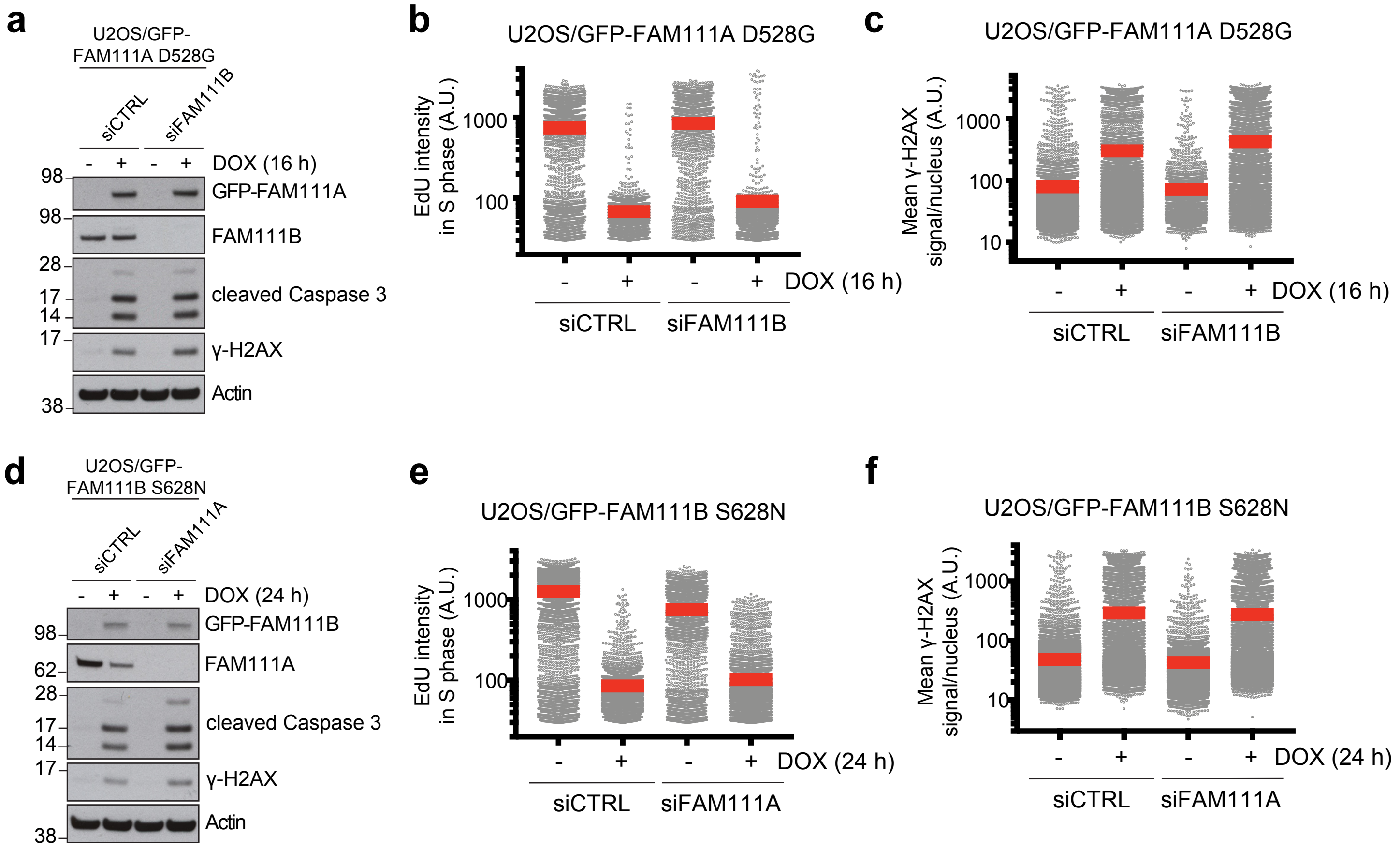
a
U2OS/GFP-FAM111A D528G
siCTRL
siFAM111B
DOX (16 h)
GFP-FAM111A
FAM111B
cleaved Caspase 3
γ-H2AX
Actin
b
U2OS/GFP-FAM111A D528G
EdU intensity in S phase (A.U.)
DOX (16 h)
siCTRL
siFAM111B
c
U2OS/GFP-FAM111A D528G
Mean γ-H2AX signal/nucleus (A.U.)
DOX (16 h)
d
U2OS/GFP-FAM111B S628N
siCTRL
siFAM111A
DOX (24 h)
GFP-FAM111B
FAM111A
cleaved Caspase 3
γ-H2AX
Actin
e
U2OS/GFP-FAM111B S628N
EdU intensity in S phase (A.U.)
DOX (24 h)
f
U2OS/GFP-FAM111B S628N
Mean γ-H2AX signal/nucleus (A.U.)
DOX (24 h)

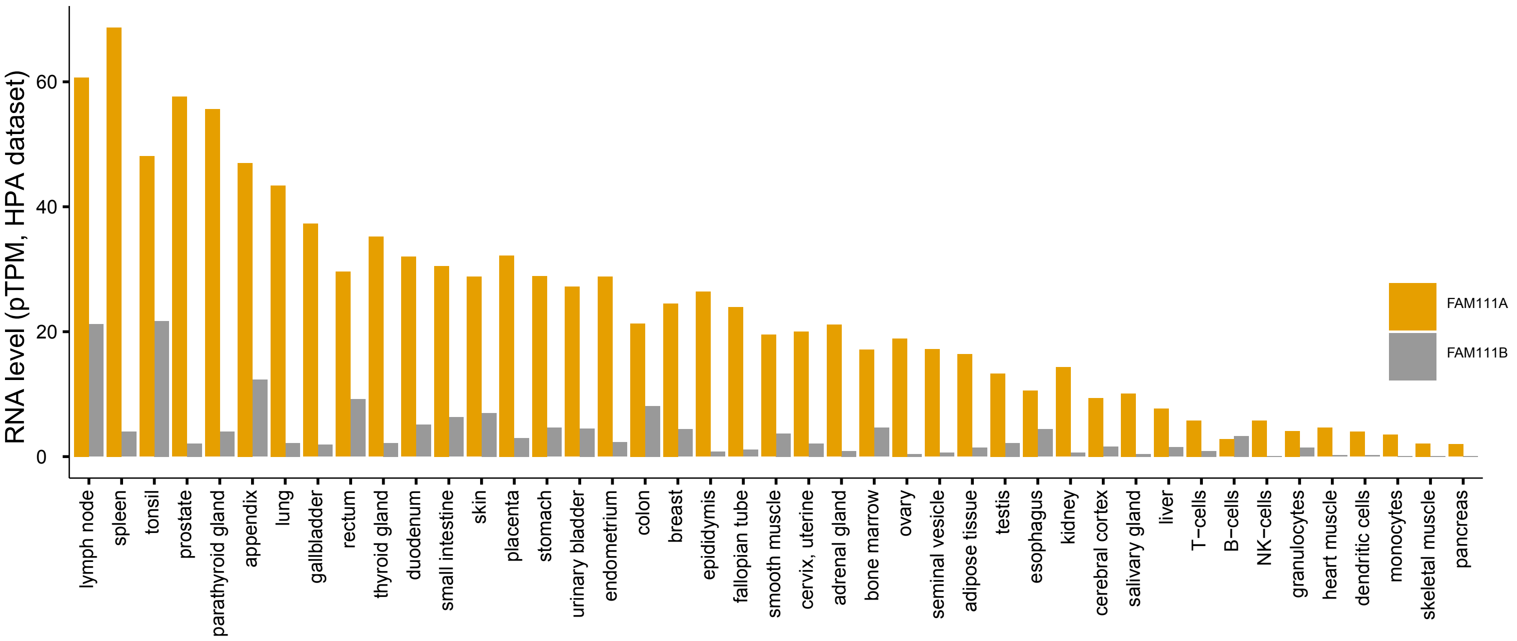
RNA level (pTPM, HPA dataset)
0
20
40
60
FAM111A
FAM111B
lymph node
spleen
tonsil
prostate
parathyroid gland
appendix
lung
gallbladder
rectum
thyroid gland
duodenum
small intestine
skin
placenta
stomach
urinary bladder
endometrium
colon
breast
epididymis
fallopian tube
smooth muscle
cervix, uterine
adrenal gland
bone marrow
ovary
seminal vesicle
adipose tissue
testis
esophagus
kidney
cerebral cortex
salivary gland
liver
T−cells
B−cells
NK−cells
granulocytes
heart muscle
dendritic cells
monocytes
skeletal muscle
pancreas
